# From individual movement to ecosystem response: how landscape structure shapes herbivore impacts

**DOI:** 10.64898/2026.11.19.766513

**Authors:** Daniela Lioren Suárez, Nara Guisoni, María Fabiana Laguna

## Abstract

Understanding how local interactions generate large-scale ecological dynamics is a central challenge in ecology. In heterogeneous environments, consumer movement links resource distribution with exploitation, producing responses that may not be predicted from consumer abundance or resource availability alone.

Here, we develop a spatially explicit stochastic individual-based model of two herbivore populations with contrasting movement and foraging strategies that share renewable resources. The model incorporates heterogeneous resources, consumer movement and feeding, resource regeneration, predation, and refuge availability.

Using Monte Carlo simulations, we examine how herbivore pressure and landscape structure shape population dynamics and resource depletion. Movement strongly determines the spatial pattern of depletion, while resource vulnerability depends on the overlap between resource distribution and consumer activity. Counterintuitively, increasing the extent of a productive region increases initial resource biomass but can also increase relative resource loss. Additional resources sustain consumer activity where herbivores concentrate, showing that greater resource availability does not necessarily buffer ecosystems against herbivory. Resource biomass can also decline substantially before major reductions in vegetation occupancy become apparent.

These results demonstrate how individual behaviour and landscape structure jointly shape ecosystem responses. The model provides a framework for investigating how spatially coupled consumer–resource interactions generate ecological dynamics.

## 1 Introduction

Consumer-resource interactions are fundamental drivers of ecological dynamics across multiple spatial and temporal scales. Resource availability influences consumer movement and feeding activity, while consumption alters resource abundance and can affect how resources are distributed across the landscape over time. Understanding how these interactions generate ecosystem-level patterns from individual-level processes remains a central challenge in ecology.

Herbivory provides a particularly important example of these processes. Through the consumption of plant resources, herbivores influence vegetation dynamics, nutrient cycling, species coexistence and ecosystem functioning across a wide range of ecosystems [1, 2, 3]. By changing resource availability, herbivores affect their own subsequent feeding activity as well as the resources available to other consumers, linking local patterns of resource consumption to changes in vegetation and herbivore distribution across the landscape.

Spatial heterogeneity is a fundamental characteristic of ecological systems and strongly influences consumer-resource interactions. Environmental gradients and variation in habitat conditions generate heterogeneous distributions of resources, which in turn affect consumer movement, resource use and competitive interactions [4, 5, 6]. When multiple consumers exploit shared resources, the effects of competition may depend on how resources are distributed across the landscape, influencing the extent and location of resource depletion and the resulting dynamics of the system [7].

These challenges become even greater when consumers differ substantially in their ecological strategies. Differences in mobility, resource preferences, consumption rates and responses to resource availability can lead to nonlinear dynamics and emergent spatial patterns through feedbacks between consumer behavior and resource distribution [8, 9]. Consumers may also modify habitat features that indirectly affect other species, for example by altering the availability of refuges from predators. Such non-trophic interactions introduce additional feedbacks between resource use, habitat structure and population dynamics. Understanding these interacting mechanisms is particularly relevant in ecosystems where consumers with different movement and foraging strategies exploit shared resources.

Patagonian wetlands, locally known as mallines, provide an illustrative example of such systems. These highly productive wetlands are embedded within the surrounding semi-arid steppe and exhibit marked spatial variation in vegetation composition and productivity [10]. This heterogeneity results in a spatially variable distribution of forage resources exploited by both native herbivores and livestock. Because these herbivores may share plant resources, their interactions can affect vegetation dynamics and the availability of forage in grazing systems [11].

Herbivorous insects also contribute to these interactions. Grasshoppers, in particular, are abundant in these ecosystems and may substantially affect forage availability during outbreak events [12, 13]. Previous studies have shown that these species exhibit selective feeding behavior associated with plant nutritional traits [14, 15]. As several of their preferred plant species are also consumed by livestock [16], these observations suggest the existence of indirect interactions mediated by shared renewable resources. Beyond these trophic interactions, shrubs provide refuge for grasshoppers against aerial predators but may also be consumed by livestock. Livestock browsing could therefore alter shrub biomass and refuge availability, potentially modifying predation risk and introducing an additional pathway through which grazing may influence grasshopper dynamics. Together, these processes illustrate how trophic and non-trophic interactions may jointly shape herbivore dynamics in heterogeneous landscapes.

Mathematical modeling provides a framework for linking individual-level ecological mechanisms to emergent ecosystem dynamics. In particular, stochastic individual-based models are well suited to heterogeneous environments because they explicitly represent movement, resource consumption, mortality and local interactions while preserving spatial structure [17, 9, 18]. They can therefore help identify mechanisms underlying ecological patterns that are difficult to isolate experimentally.

In a previous study, our research group developed a spatially explicit stochastic individual-based model inspired by Patagonian wetlands to investigate the impact of herbivorous insects on resource dynamics [19]. The model incorporated heterogeneous resource distributions, species-specific feeding preferences and movement guided by resource availability.

It was used to examine how grasshopper density affects vegetation depletion and to identify conditions under which outbreaks may substantially reduce resource availability.

Building on that approach, the present work investigates how herbivores with contrasting movement and foraging strategies interact through shared renewable resources in a heterogeneous landscape. The model incorporates consumer-specific movement behaviors and resource preferences, resource consumption and regeneration, and predation risk associated with habitat structure, allowing us to examine how the spatial relationship between consumers and resources affects ecosystem responses to increasing herbivore abundance.

Using Monte Carlo simulations, we analyze how consumer density, resource availability and landscape heterogeneity influence population dynamics and resource depletion. In particular, we ask whether increasing resource availability necessarily reduces the impact of herbivory, or whether the outcome depends on the spatial relationship between resources and consumer activity. By explicitly linking individual movement and feeding behavior to resource dynamics and mortality, this framework provides a way to investigate how interactions between consumers and their heterogeneous environment generate ecosystem-level patterns.

## 2 Methods

We develop a spatially explicit stochastic individual-based model describing the interactions between two herbivore populations with contrasting ecological strategies and renewable resources distributed across a heterogeneous landscape. The framework is motivated by ecological processes observed in Patagonian wetlands and builds upon previous work on grasshopper-resource dynamics [19]. Empirical studies of herbivore populations, grazing systems and vegetation in Patagonian wetlands provided the basis for selecting and parameterizing model processes and values [19, 14, 20]. The model explicitly represents consumer movement, resource consumption, resource regeneration, predation and refuge effects. A detailed description of the model formulation, parameterization and implementation, together with a schematic overview of the sequence of stochastic processes and interactions, is provided in the Supplementary Material.

### Biological components

The model describes two herbivore populations, *H*_1_ and *H*_2_, interacting with three renewable vegetation resources, *R*_1_, *R*_2_ and *R*_3_, and shrub cover, *R*_4_. The two herbivores differ in movement strategies and resource use. The first population, *H*_1_, represents a mobile herbivore whose movement is biased toward more productive regions and whose feeding is resource selective. The second population, *H*_2_, represents a larger herbivore that moves without directional bias, consumes the same renewable resources as *H*_1_ and additionally browses on shrubs. Shrubs provide refuge for *H*_1_ by reducing predation risk and therefore mediate an indirect interaction between the two herbivore populations.

The three renewable resources differ in abundance, productivity, spatial distribution and attractiveness to consumers. Their biomass changes through consumption and regeneration, whereas shrub biomass does not regenerate over the time scale considered. For each renewable resource, biomass is bounded by a maximum value determined by the local carrying capacity. The parameter *R* denotes the combined carrying capacity of the resources per patch and represents local resource availability. The initial total biomass across the landscape depends on the resource composition and spatial distribution, as well as on the landscape structure. The two herbivore populations also differ in their resource preferences: *H*_1_ prefers *R*_1_ *> R*_2_ *> R*_3_, whereas *H*_2_ shows equal preference for *R*_1_ and *R*_2_, followed by *R*_3_ and *R*_4_.

Herbivore reproduction is not included because the model is designed to represent a timescale corresponding approximately to one generation of the faster-reproducing population. The abundance of *H*_1_ can decrease through mortality, whereas the abundance of *H*_2_ remains constant throughout each simulation because its characteristic timescale is longer than the simulated period. In addition, both herbivore populations are subject to local carrying capacities.

### Landscape and initial conditions

The landscape is represented by a two-dimensional lattice of *L*_*x*_ × *L*_*y*_ patches. Each patch may contain herbivores, renewable resources and shrubs.

To represent environmental heterogeneity, the landscape is divided into a higher-productivity region A and a lower-productivity region B (Fig. 1 A). Resource abundance and composition differ between these regions. Shrubs occur in region B and provide refuge for *H*_1_, while also constituting a secondary food resource for *H*_2_. The model uses a square lattice for the simulations presented here, although the formulation is not restricted to this geometry.

**Figure 1:**
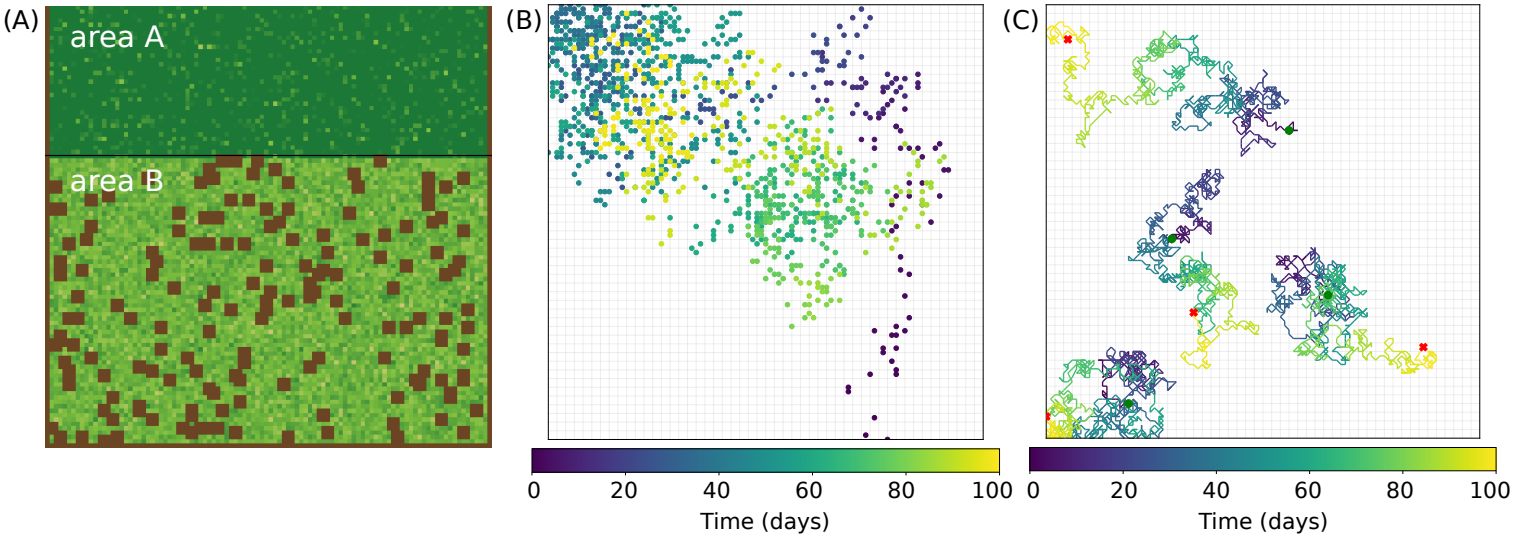
Landscape structure and individual movement in the model. (A) Simplified representation of the landscape showing areas A and B. Brown patches represent bare soil, and the darker green shading in area A represents greater resource abundance. (B) Trajectory of a representative *H*_1_ individual, showing its progressive movement from area B toward area A. (C) Trajectories of four representative *H*_2_ individuals showing spatially unbiased movement across the landscape. Green points indicate the beginnings of the random walks, while red crosses indicate their endpoints.

Initial resource distributions are prescribed according to the spatial heterogeneity of the landscape. Resources *R*_1_ and *R*_3_ are distributed between regions A and B, whereas *R*_2_ is preferentially distributed in region A. Shrubs are restricted to region B, which also contains bare-soil patches. The two herbivore populations are initialized independently. Individuals of *H*_1_ are initially placed along selected landscape boundaries, representing immigration from the surrounding landscape, whereas *H*_2_ individuals are distributed throughout the landscape. The initial abundances of both herbivores are varied among simulations. The initial spatial distributions of resources and herbivores are illustrated in Fig. S2A of the Supplementary Material.

### Consumer movement

The two herbivore populations follow different movement strategies, illustrated in Fig. 1 B,C. Individuals of *H*_1_ perform directional movement biased toward the higher-resource region A. Movement distances are variable, with the probability of long-range displacements decreasing with distance. Once within region A, movement becomes unbiased. In contrast, *H*_2_ performs an unbiased random walk among neighboring patches. Migration is constrained by local carrying capacity and resource availability. The complete movement rules and parameterization are given in the Supplementary Material.

### Resource consumption and regeneration

Both herbivore populations consume *R*_1_, *R*_2_ and *R*_3_ and therefore interact indirectly through resource depletion. Consumption depends on consumer-specific feeding rates and resource preferences. In the parameterization considered here, both populations preferentially consume some resource types over others, while *H*_2_ additionally consumes shrub vegetation (*R*_4_).

Renewable resources regenerate through a density-dependent process after consumption. Resource types may differ in their regeneration rates and spatial distributions. In contrast, shrub biomass does not regenerate during the simulations, so browsing by *H*_2_ progressively reduces refuge availability for *H*_1_. The detailed consumption and regeneration rules are provided in the Supplementary Material.

### Mortality

Individuals of *H*_1_ are subject to predation, with local predation risk decreasing with shrub cover. They may also experience mortality through interactions with *H*_2_, representing processes such as trampling or incidental mortality during grazing. Thus, *H*_2_ affects *H*_1_ through both competition for shared resources and habitat modification.

Mortality is activated when the total local vegetation cover, *R*_*T*_ = *R*_1_ + *R*_2_ + *R*_3_ + *R*_4_, falls below a threshold *R*^*^. Under this condition, an *H*_1_ individual may die through predation or through interaction with *H*_2_. The complete stochastic rules are given in the Supplementary Material.

### Time evolution

The model evolves through stochastic elementary events corresponding to resource consumption, mortality or migration. For each update attempt, a landscape patch is selected and one of these processes is chosen according to prescribed probabilities. One Monte Carlo step (MCS) consists of *N* = *L*_*x*_ × *L*_*y*_ elementary update attempts. The correspon-dence between MCS and biological time is established using a known biological timescale, such as the time required for *H*_1_ to reach the region with higher resource availability. We consider a simulation period of 100 biological days, corresponding to 10^4^ MCS.

### Simulation experiments

We used Monte Carlo simulations to investigate how consumer abundance, resource availability and landscape structure affect the emergent dynamics of the system. Unless otherwise stated, model parameters were fixed at the values reported in Table S1. We varied the initial number of *H*_1_ individuals (0 - 150,000), the number of *H*_2_ individuals (0 - 100), and the resource carrying capacity *R* (400 - 1000 g m^−2^), together with the landscape conditions specified for each experiment. The landscape consisted of *L*_*x*_ = *L*_*y*_ = 100 patches, each representing an area of 1 m^2^.

For each parameter combination, the number of independent realizations is specified for each case. We monitored consumer abundance, resource biomass, their spatial distributions and ecosystem-level dynamics. Full parameter values and simulation settings are provided in Table S1 and the Supplementary Material.

## 3 Results

The results are organized in three parts. We first illustrate the behavior emerging from the model under a representative set of parameters to introduce the main ecological mechanisms. We then investigate how the ecosystem responds to increasing herbivore pressure. Finally, we examine how the relative extent of the productive and less productive regions influences resource depletion and ecosystem responses to herbivory.

### 3.1 Emergent dynamics in a heterogeneous landscape

We first examine how individual movement rules translate into population- and ecosystem-level spatial patterns. Panels B and C of Fig. 1 illustrate the contrasting movement strategies of the two herbivore populations. Individuals of *H*_1_ enter the landscape from the boundaries of area B and progressively move toward the resource-rich region A, where their movement becomes more localized. Their trajectories also exhibit long-range displacements, reflecting their ability to move over larger distances. In contrast, *H*_2_ individuals perform an unbiased random walk among neighboring patches, resulting in more spatially dispersed trajectories across the landscape.

These contrasting movement strategies generate distinct spatial patterns of herbivory and provide the basis for the subsequent ecosystem-level dynamics. Figure 2A shows two representative stages of a simulation with 30 individuals of *H*_2_ and an initial *H*_1_ density of 9 individuals m^−2^. The spatial snapshots illustrate the early stages of the simulation and the final state after 100 biological days, showing the progressive entry and redistribution of *H*_1_ and the associated changes in vegetation. Because *H*_1_ individuals enter progressively from the landscape boundaries, the initial stages do not yet reflect the full spatial distribution of the population assigned to the simulation.

**Figure 2:**
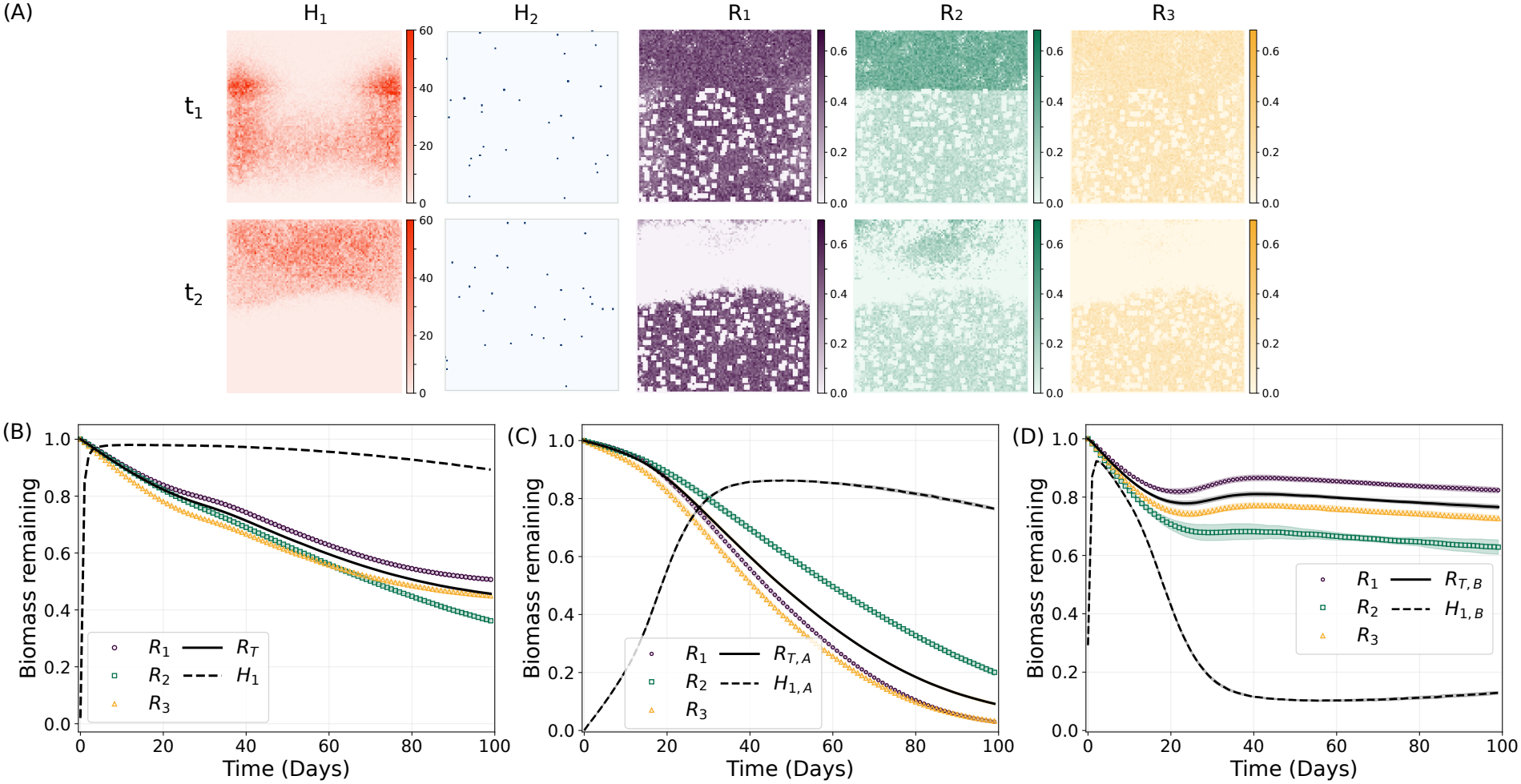
Emergent dynamics in a heterogeneous landscape for a representative simulation with an initial *H*_1_ density of 9 individuals m^−2^, 30 individuals of *H*_2_, and *R* = 700 g m^−2^. (A) Spatial distributions of *H*_1_, *H*_2_, *R*_1_, *R*_2_, and *R*_3_ at days *t*_1_ = 10 and *t*_2_ = 100. (B) Temporal evolution of the normalized biomass of each renewable resource, total renewable biomass (*R*_T_), and the *H*_1_ population. (C–D) Same quantities as in (B), restricted to areas A and B, respectively. Panels (B–D) show averages over five independent simulations, with shaded areas indicating standard deviations. Resource biomass is normalized by the initial total biomass of each resource, and the *H*_1_ population by its initial total value.

During the initial stages, *H*_1_ individuals enter through area B, the less productive region, and move toward area A. By the end of the simulation, most *H*_1_ individuals have reached and become concentrated in area A, where their grazing activity reduces vegetation biomass. Movement within area A persists, although individuals may occasionally return to area B as resources become depleted in area A. In contrast, the more homogeneous distribution of *H*_2_ generates a more spatially diffuse grazing pressure. The Supplementary Material (Fig. S2) shows the spatial distributions at additional stages throughout the simulation.

The temporal dynamics are shown in panels B-D of Fig. 2. Panel B shows the normalized biomass of each renewable resource, total renewable biomass (*R*_T_) and the normalized abundance of *H*_1_. As *H*_1_ individuals enter the landscape, they become subject to the mortality processes described above, while those still outside continue to enter progressively. Consequently, the number of *H*_1_ individuals within the landscape initially increases as the population enters and moves through the system. Over the same period, the biomass of renewable resources decreases, although the magnitude of the reduction differs among resources.

Panels C and D show resource biomass and *H*_1_ abundance in areas A and B, respectively, illustrating how the movement of *H*_1_ between the two regions shapes their local dynamics. In area B (Fig. 2D), resource biomass initially decreases as *H*_1_ enters and grazes this region. As *H*_1_ individuals move toward area A, grazing pressure in B declines and resource biomass shows a slight increase. In area A (Fig. 2C), by contrast, resource biomass decreases as *H*_1_ progressively concentrates there. At later times, depletion of resources in A is followed by the return of *H*_1_ to B, where renewed grazing leads to a further reduction in resource biomass. The curves for areas A and B thus show how the movement of *H*_1_ produces a sequence of resource depletion and recovery in the two regions.

The simulations also reveal a difference between changes in resource biomass and changes in vegetation occupancy, which is shown in Supplementary Material, Fig. S3. While resource biomass decreases continuously as vegetation is consumed, a patch remains classified as vegetated until its entire vegetation biomass has been depleted. Consequently, substantial biomass losses can occur before the fraction of vegetated patches decreases. Vegetation occupancy therefore provides a more discrete measure of vegetation loss, whereas biomass captures changes in resource availability within patches that remain vegetated.

Together, these results show how individual movement rules generate spatial redistribution of herbivores and, through their interactions with heterogeneous resources, produce emergent ecosystem-level dynamics.

### 3.2 Ecosystem response to herbivore pressure

Having characterized the mechanisms responsible for the emergent spatial organization of the system, we next examine how vegetation responds to increasing herbivore pressure. We consider the effects of the two herbivore populations separately, varying the abundance of one population while keeping the other fixed.

Figure 3A shows the biomass remaining after 100 days for each renewable resource as a function of initial *H*_1_ density, with the abundance of *H*_2_ fixed at 30 individuals and a carrying capacity of 700 g m^−2^. The biomass of all three resources decreases gradually as *H*_1_ density increases, but the magnitude of the response differs among resources. *R*_2_ shows the strongest reduction, whereas *R*_1_ and *R*_3_ retain larger fractions of their initial biomass. The response of individual resources to increasing *H*_1_ density for different values of *R* is further examined in the Supplementary Material (Fig. S4).

**Figure 3:**
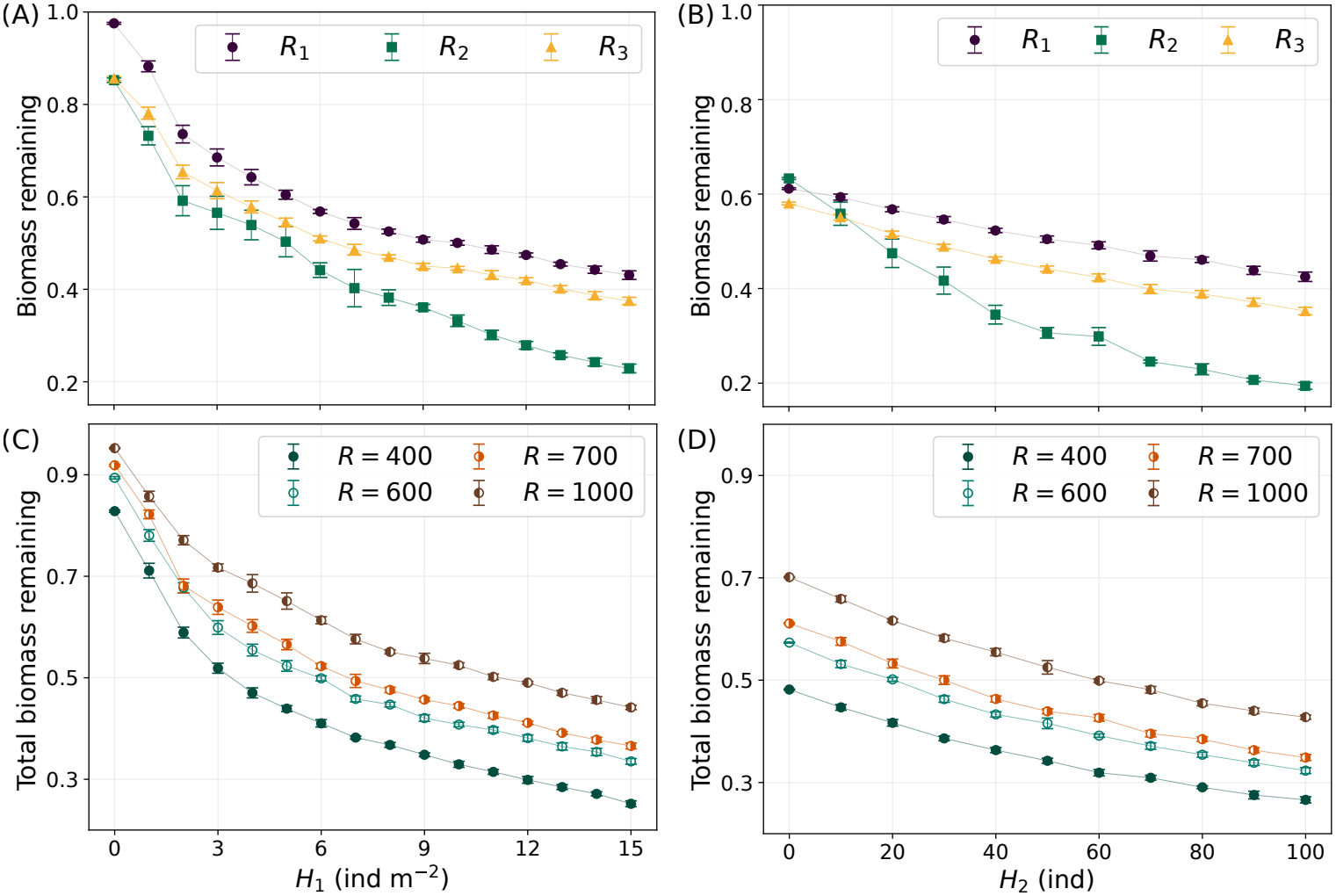
Response of vegetation biomass to increasing herbivore pressure. (A) Biomass remaining after 100 days for each renewable resource as a function of initial *H*_1_ density, for a fixed abundance of 30 individuals of *H*_2_ and *R* = 700 g m^−2^. (B) Corresponding response to increasing *H*_2_ abundance, for a fixed initial *H*_1_ density of 7 individuals m^−2^ and *R* = 700 g m^−2^. In both cases biomass is normalized by the initial biomass of the corresponding resource. (C) Total biomass remaining after 100 days as a function of initial *H*_1_ density and *H*_2_ = 30, for four values of *R* (400, 600, 700 and 1000 g m^−2^). Total biomass is calculated from the sum of the biomasses of the three renewable resources and normalized by the corresponding initial total renewable-resource biomass. (D) Corresponding total biomass remaining as a function of *H*_2_ abundance and a initial density of *H*_1_ =7 individuals per *m*^−2^ for the same four values of *R*. Each data point represents the final biomass, averaged over the last 100 MCS of each of five independent runs. Error bars indicate the corresponding standard deviations.

Figure 3B shows the response to increasing *H*_2_ abundance, with the initial density of *H*_1_ fixed at 7 individuals m^−2^ and the carrying capacity also fixed at 700 g m^−2^. Again, the biomass of all three resources decreases gradually with increasing herbivore abundance. Interestingly, the relative ranking of the resources changes as *H*_2_ abundance increases. At low *H*_2_ abundance, *R*_2_ retains the largest fraction of its initial biomass, but its biomass decreases more rapidly than that of the other resources and becomes the lowest at higher *H*_2_ abundances. Thus, the resource that is initially least affected under this condition becomes the most strongly affected as herbivore pressure increases. This change in relative response suggests that resource vulnerability depends not only on initial biomass, but also on the interaction between spatial distribution, herbivore movement, consumption and resource regeneration.

The stronger response of *R*_2_ may result from the interaction between resource distribution and herbivore movement. Unlike *R*_1_ and *R*_3_, which are distributed across both regions, *R*_2_ is preferentially concentrated in area A, toward which *H*_1_ progressively moves and where it spends much of its time feeding. Although *R*_2_ is the second-ranked resource in *H*_1_’s preference hierarchy, its spatial concentration exposes it to sustained grazing pressure as *H*_1_ accumulates in area A. Increasing *H*_1_ abundance further intensifies this pressure. The response to *H*_2_ is more spatially homogeneous because of its unbiased movement, although the concentration of *R*_2_ in area A may still contribute to its greater reduction. Moreover, *R*_2_ regenerates more slowly than *R*_1_ and *R*_3_, limiting its capacity to recover from grazing and potentially amplifying its biomass reduction over time. Together, resource distribution, herbivore movement, resource preferences, and regeneration rates may explain the contrasting responses among resource types.

Panels C and D show the corresponding total biomass remaining after 100 days, calculated as the sum of *R*_1_, *R*_2_ and *R*_3_, and normalized by their combined initial biomass, for different values of resource carrying capacity *R*. Total biomass decreases with increasing abundance of either herbivore, while the initial vegetation biomass modulates the fraction that remains after 100 days. The aggregate response therefore reflects the combined effects of herbivore abundance, spatial resource distribution and resource dynamics.

Together, these results show that the response of vegetation to herbivore pressure depends not only on the abundance of consumers but also on the spatial arrangement and dynamics of the resources.

### 3.3 Landscape structure modulates ecosystem response to herbivore pressure

The results above show that vegetation responses to herbivore pressure depend not only on herbivore abundance but also on landscape structure. We therefore examined how the relative extent of the productive and less productive regions affects vegetation biomass, keeping herbivore abundances and the resource biomass assigned to patches in each region unchanged. We varied the ratio *f*_*A*_*/f*_*B*_, where *f*_*A*_ and *f*_*B*_ = 1 − *f*_*A*_ denote the fractions of the landscape occupied by regions A and B, respectively. Increasing the extent of region A therefore increases the total initial resource biomass across the landscape without changing the local carrying capacity *R* (see Supplementary Material, Fig. S5).

Figure 4A shows the fraction of initial biomass remaining after 100 days for each of the three renewable resources as a function of *f*_*A*_*/f*_*B*_. For all three resources, this fraction decreases as the relative extent of region A increases, although the magnitude of the response differs among resources. Thus, increasing the total initial resource biomass does not reduce the relative impact of herbivory. This response may be related to the movement strategy of *H*_1_, whose individuals progressively move toward region A, where movement subsequently becomes more localized. Expanding region A therefore increases the amount of resource available in the region where *H*_1_ concentrates its activity.

**Figure 4:**
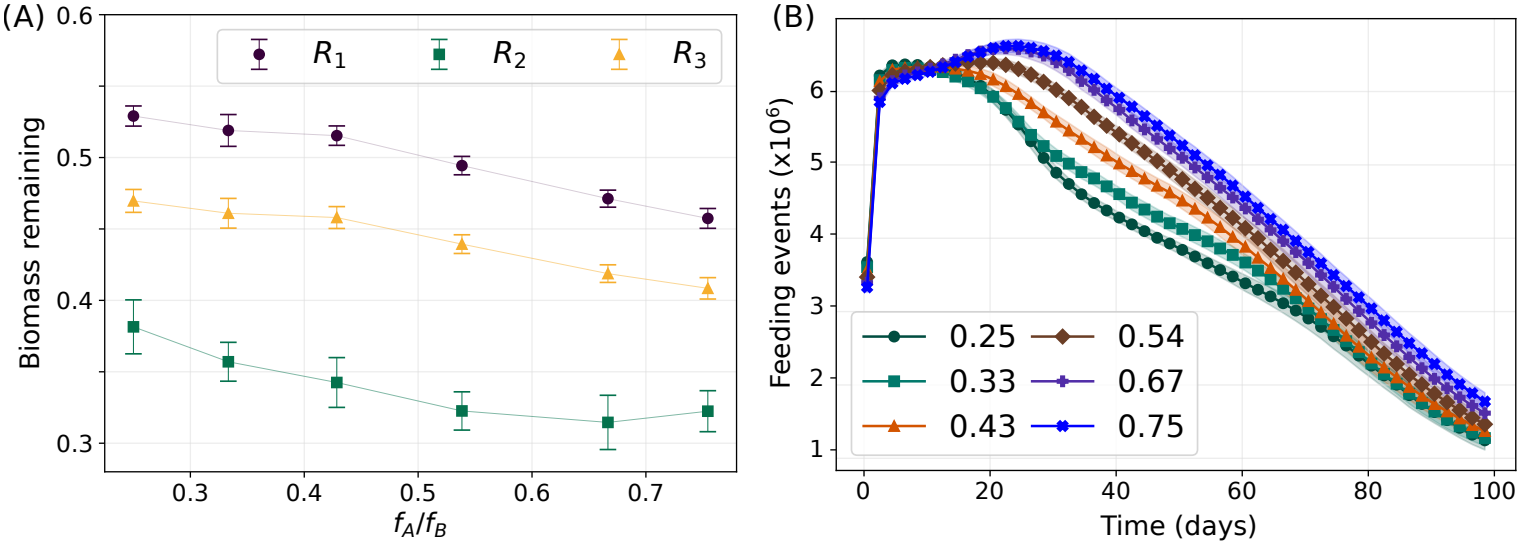
(A) Fraction of biomass remaining after 100 days as a function of the relative size of the productive regions, *f*_*A*_*/f*_*B*_, with *H*_1_ fixed at 10 individuals m^−2^, *H*_2_ fixed at 30 individuals, and *R* = 700 g m^−2^. (B) Temporal evolution of the number of feeding events for *H*_1_ and *H*_2_ for the different values of *f*_*A*_*/f*_*B*_ shown in (A) and indicated in the panel.

To investigate how feeding activity varies across landscape configurations, we analyzed the temporal dynamics of feeding events. Figure 4B shows the daily total number of successful feeding events for the different values of *f*_*A*_*/f*_*B*_. During the initial stage of the simulations, feeding activity is similar across landscape configurations. Differences emerge later, with larger values of *f*_*A*_*/f*_*B*_ sustaining higher feeding activity, particularly during the intermediate stages of the simulation.

The resource-specific dynamics in Supplementary Material, Fig. S6, show that this increase is most pronounced for *R*_2_, which is preferentially distributed in region A. By contrast, feeding activity involving *R*_1_ and *R*_3_ varies less across landscape configurations. This pattern is consistent with the spatial overlap between *R*_2_ and the region toward which *H*_1_ moves, where its activity subsequently becomes more localized.

The initial and final biomass of *R*_2_ provide complementary evidence for this interpretation (Supplementary Material, Fig. S5). Although its initial biomass increases with the extent of region A, its final biomass remains approximately constant. The additional initial availability therefore does not translate into a greater final biomass, consistent with increased net resource depletion in landscapes with a larger productive region. This result, together with the feeding dynamics, suggests that expanding region A sustains greater resource use rather than simply buffering vegetation against herbivore pressure.

Overall, these results show that the effect of landscape structure depends on how resource distribution interacts with consumer movement. Increasing the extent of a productive region can increase total resource availability while also increasing the availability of resources in the area where consumer activity is concentrated. Consequently, greater initial resource availability does not necessarily reduce the relative impact of herbivory.

## 4 Discussion

In this study, we developed a spatially explicit stochastic individual-based model to explore how herbivore populations with contrasting ecological strategies interact with renewable resources in a heterogeneous landscape. Inspired by ecological processes observed in Patagonian wetlands, the model was formulated as a general framework rather than as a detailed representation of a particular ecosystem. Its main purpose is to link individual movement, resource use, regeneration and mortality with population- and ecosystem-level dynamics, and to examine how herbivore pressure and landscape structure shape ecosystem responses.

A previous model of grasshopper impacts on Patagonian wetlands, developed by one of the authors, provides a direct precedent for this approach [19]. That model focused on a specific system, whereas the present framework represents multiple resources and two herbivore populations with contrasting movement strategies. It also explicitly considers the spatial redistribution of consumers, their interaction through shared resources and the effects of shrub availability on mortality risk. This broader formulation allows us to examine how consumer abundance and the spatial relationship between consumers and resources shape ecosystem responses, rather than focusing on degradation thresholds for a particular system.

A central result is that herbivore movement strongly influences the spatial pattern of resource depletion. *H*_1_ progressively moves toward the resource-rich region A, whereas *H*_2_ moves without directional bias and becomes more broadly distributed across the landscape. The concentration of *H*_1_ in region A produces localized resource depletion, followed by redistribution as resources become less abundant. Thus, the effect of a given herbivore abundance depends not only on the number of consumers, but also on where they concentrate their activity. This result highlights the importance of explicitly representing movement when assessing herbivore impacts in spatially heterogeneous systems.

The resource-specific responses further illustrate the importance of spatial coupling between consumers and resources. *R*_2_, which is preferentially distributed in region A, shows the strongest reduction as herbivore pressure increases. Its vulnerability reflects the spatial overlap between resource availability and herbivore activity, together with differences in consumption preferences and regeneration rates. Under increasing *H*_2_ abundance, *R*_2_ initially retains a relatively large fraction of its biomass but subsequently becomes the most strongly affected resource. Thus, resource vulnerability cannot be inferred from initial abundance alone, but depends on the interaction between resource distribution, consumer movement, consumption and regeneration.

Our results also reveal a counterintuitive effect of landscape structure. Increasing the relative extent of the productive region A increases the total initial amount of vegetation but leads to a lower fraction of biomass remaining after 100 days. This pattern occurs for all three renewable resources. A larger productive area therefore does not necessarily reduce the relative impact of herbivory.

This response can be understood from the movement strategy of *H*_1_. Because A is the region toward which *H*_1_ progressively moves and where its movement subsequently becomes more localized, increasing its extent also increases the amount of resource available within the region where herbivore activity is concentrated. The additional resource can sustain herbivore activity for longer periods and reduce the limitation imposed by local resource depletion. Thus, increasing the extent of region A does not simply dilute herbivore pressure over a larger resource pool, but may maintain access to resources in the area most strongly used by *H*_1_. The temporal dynamics of feeding activity support this interpretation, as increasing the relative extent of region A leads to higher feeding activity during the later stages of the simulation, particularly for resources preferentially distributed in A.

Resource *R*_2_ provides a clear example of this mechanism. Because it is preferentially distributed in A, increasing the extent of this region increases both its initial amount and the area over which it is accessible to *H*_1_. The greater availability of *R*_2_ therefore does not compensate for its increased exposure to herbivory; instead, the spatial overlap between resource distribution and consumer movement appears to increase the pressure exerted on this resource. More generally, the ecological effect of increasing resource availability depends not only on how much additional vegetation is present, but also on where it is located relative to consumer activity.

The dependence of herbivore impacts on initial vegetation biomass is consistent with the previous grasshopper model [19], in which the herbivore density associated with substantial biomass loss depended on the initial biomass available. The present results extend this perspective by showing that total resource abundance alone is not sufficient to characterize ecosystem response: its spatial distribution relative to consumer movement also matters. Consequently, landscapes with similar total vegetation biomass may experience different levels of resource depletion depending on how resources are distributed relative to areas of concentrated consumer activity.

Another result shared with the previous model is the distinction between vegetation biomass and vegetation occupancy. In the present simulations, biomass decreases before a substantial loss of occupied patches becomes apparent. Vegetation can therefore remain present across a large fraction of the landscape while the amount of forage available within occupied patches has already been substantially reduced. Measures based only on vegetation presence or cover may consequently underestimate early effects of herbivory, whereas biomass can provide a more sensitive indicator of declining resource availability. The recurrence of this pattern in models with different levels of ecological detail suggests that it may reflect a general consequence of progressive resource depletion within occupied patches.

Taken together, these results show that herbivore impacts emerge from the interaction between consumer abundance, resource availability and landscape structure. Consumer abundance alone is therefore insufficient to characterize ecosystem response to herbivory. The same number of consumers may produce different outcomes depending on the amount of vegetation available, its spatial distribution and its overlap with consumer activity. Landscape structure is thus not simply a background condition, but an active component of consumer-resource interactions.

The model is intentionally formulated as a simplified representation of these processes. It does not explicitly represent reproduction and includes a limited number of resource types and simplified movement and mortality rules. Its purpose is not to reproduce the dynamics of a particular wetland in all their detail, but to identify mechanisms through which individual behavior and landscape structure can influence ecosystem-level responses. The framework can also be adapted to other consumer-resource systems by modifying resource types, consumer strategies, demographic processes, or parameter values.

Overall, our results indicate that ecosystem response to herbivore pressure depends jointly on consumer abundance, initial resource availability and the spatial relationship between consumers and resources. In particular, greater resource availability does not necessarily reduce the impact of herbivory when additional resources are located in areas where consumers concentrate their activity. Considering both the amount of vegetation available and its spatial organization relative to consumer movement therefore provides a more complete picture of ecosystem response than either factor considered in isolation. More broadly, our results highlight that understanding herbivore impacts requires considering not only how much vegetation and how many consumers are present, but also how their spatial distributions interact over time.

## Acknowledgments

We thank L. S. Serrano and V. Fernández-Arhex for fruitful discussions and valuable insights that contributed to this work.

## Notes

### Competing Interest Statement

The authors have declared no competing interest.

