## Supplementary Material for "From individual movement to ecosystem response: how landscape structure shapes herbivore impacts"

Daniela Lioren Suárez, Nara Guisoni and María Fabiana Laguna

#### 1 Supplementary Methods

##### Model overview

The model describes the spatial and temporal dynamics of two herbivore populations, $H_1$ and $H_2$, interacting with renewable vegetation resources in a heterogeneous landscape. The two populations differ in their movement strategies and resource use. Individuals of $H_1$ progressively enter the landscape from its boundaries and are biased toward the resource-rich region A, whereas $H_2$ individuals move through the landscape without a directional bias. Both populations consume the same renewable resources, while $H_2$ also consumes shrub biomass, which provides refuge for $H_1$.

The model evolves through a sequence of stochastic elementary events involving herbivore movement, resource consumption, resource regeneration and mortality. At each update attempt, a landscape patch is selected and one of these processes is attempted according to the corresponding event probabilities. Movement changes the spatial distribution of the herbivores, consumption reduces local resource biomass and regeneration restores renewable resources. Mortality affects $H_1$ through two mechanisms: predation, whose risk depends on local vegetation cover and is reduced by the availability of shrubs as refuge, and direct interactions with $H_2$, which may cause incidental mortality during grazing or through trampling.

##### Model implementation

This section provides the detailed formulation and implementation of the model described in the main text. The complete parameterization used in the simulations is reported in Table S1.

###### Landscape and initial conditions

The landscape consists of a square lattice of $L_x \times L_y$ patches. In the simulations presented here, $L_x = L_y = 100$, resulting in $N = 10,000$ patches. The landscape is divided into two regions, A and B, with different levels of resource availability. The fraction of the landscape occupied by each region is specified by $f_A$ and $f_B = 1 - f_A$.

The initial distribution of renewable resources is prescribed according to the spatial structure of the landscape. Resources $R_1$ and $R_3$ are distributed randomly between regions A and B, whereas $R_2$ is preferentially distributed in region A. Shrubs ($R_4$) occur exclusively in region B, which also contains patches of bare soil where vegetation is absent. The local carrying capacity $R$ defines the maximum combined biomass of renewable resources and shrubs in a vegetated patch $(i, j)$ and serves as a measure of landscape productivity. The actual vegetation biomass in a patch may be lower than $R$, depending on its resource composition and initial state, and is zero in bare-soil patches. Thus, the total vegetation biomass in each patch satisfies

$$R_1(i, j) + R_2(i, j) + R_3(i, j) + R_4(i, j) \leq R. \tag{S1}$$

Consequently, the total initial vegetation biomass of the landscape depends on both the fraction of patches occupied by vegetation and their initial biomass, and is lower than $RL_xL_y$ when some patches contain less than the maximum biomass or are bare.

In the parameterization considered here, 40% of the landscape is allocated to $R_1$, 20% to $R_2$ (with 60% allocated to area A and 40% to area B), 15% to $R_3$, and 10% to shrubs. The remaining 15% consists of bare-soil patches, where all vegetation components are absent. The resource fractions define the initial composition of vegetated patches and are kept fixed unless otherwise stated. With this parameterization, the total initial vegetation biomass is $0.85RL_xL_y$, assuming that all non-bare patches initially contain biomass equal to $R$.

The initial number of $H_1$ and $H_2$ individuals are control parameters. Individuals of $H_1$ are initially placed along selected boundaries of the landscape, representing immigration from surrounding areas. Individuals of $H_2$ are distributed randomly throughout the landscape according to the prescribed initial abundance. The local carrying capacity of $H_1$ is fixed at $T = 60$ individuals per patch, whereas at most one $H_2$ individual can occupy a patch.

Reproduction is not explicitly represented because the model is designed to capture ecological dynamics over a timescale corresponding approximately to one generation of the faster-reproducing population. Over this period, changes in herbivore abundance are therefore driven by mortality rather than reproduction. Consequently, the number of $H_1$ individuals can decrease through mortality, whereas the number of $H_2$ individuals remains constant throughout each simulation, reflecting its longer characteristic timescale.

##### Movement rules

Movement is implemented as a stochastic process and differs between the two herbivore populations. Individuals of $H_1$ can move over variable distances and exhibit a directional bias relative to the higher-resource region A. The migration probability is defined as

$$M_\alpha = \frac{m_\alpha}{d}, \tag{S2}$$

where $d$ is the distance between the current patch and the candidate destination, and $m_\alpha$ is a probability that depends on the direction of movement relative to region A.

For an arbitrary patch $(i, j)$, the direction parameter $\alpha$ takes four possible values. The value of $\alpha$ depends on the direction of movement relative to region A: $\alpha = 1$ for movement toward region A, $\alpha = 2$ for movement away from region A, and $\alpha = 3$ for lateral movement that maintains the same distance from region A. For individuals already located within region A, $\alpha = 0$, corresponding to unbiased movement with respect to the direction of region A. The distance between patches is defined as

$$d = \sqrt{\Delta i^2 + \Delta j^2}, \tag{S3}$$

where $\Delta i$ and $\Delta j$ are the displacements in the two lattice directions.

Individuals of $H_2$ perform an unbiased random walk. They migrate randomly to first- and second-order neighboring patches with probability $m'$. Unlike $H_1$, their movement does not depend on the location of region A or on local resource abundance.

For both populations, migration is constrained by the carrying capacity of the destination patch, denoted by $T$ for $H_1$ and $G$ for $H_2$. In addition, when the current patch does not contain sufficient resources for feeding, migration is mandatory.

The migration process can be written schematically as

$$H_1(i, j)E_1(i', j') \xrightarrow{M_\alpha} E_1(i, j)H_1(i', j') \tag{S4}$$

$$H_2(i, j)E_2(i'', j'') \xrightarrow{m'} E_2(i, j)H_2(i'', j''), \tag{S5}$$

where $(i, j)$ denotes the current position of $H_1$ and $H_2$, respectively, $E_l$ $(l = 1, 2)$ denotes the space available within a patch for species $l$ to migrate into, and the primed coordinates denote candidate destination patches.

##### Resource consumption

Both herbivore populations consume the same renewable resources and therefore compete through resource depletion. Consumption is determined by consumer-specific feeding rates and resource preferences. For an individual of population $H_m$, the probability of consuming resource $R_n$ is denoted by $c_{m,n}$, with

$$\sum_j c_{m,n} = 1 \tag{S6}$$

where $m$ denotes the species and $n$ the resource. Equation S6 indicates that, given a feeding event, the species consumes at least one of the available resources.

In the parameterization considered here, the three renewable resources are ranked according to their preference by the consumers. For $H_1$,

$$c_{1,1} > c_{1,2} > c_{1,3}, \tag{S7}$$

and for $H_2$,

$$c_{2,1} = c_{2,2} > c_{2,3} = c_{2,4}. \tag{S8}$$

The corresponding preferences of $H_2$ and all numerical values of the consumption probabilities are given in Table S1.

The amount of biomass removed during a feeding event depends on the feeding rate of the consumer population and on the duration represented by the event. Feeding rates expressed on a daily time scale are converted to the corresponding biomass removed per Monte Carlo update using the temporal calibration described below. The parameterization of $H_2$ resource consumption is informed by field observations of sheep and goats [1]. Feeding process can be described schematically as

$$H_m(i,j)R_n(i,j) \xrightarrow{c_{m,n}} H_m(i,j)(R_n(i,j) - \Delta R_{m,n}) \tag{S9}$$

where $(i,j)$ denotes an arbitrary patch, and $\Delta R_{m,n}$ is the amount of resource $n$ consumed by species $m$ during a feeding event.

In addition to the renewable resources, $H_2$ consumes shrub biomass. Shrub consumption reduces $R_4$ and therefore progressively decreases the refuge available to $H_1$. Because shrubs do not regenerate during the simulations, this effect is cumulative.

#### Resource regeneration

Renewable resources can regenerate at regular time intervals, with the regeneration rates depending on the resource type. Regeneration depends on the resource density within each patch and on the presence of herbivores of species $H_1$. For a given resource $R_n$ to regenerate in a patch $(i,j)$, the patch must not be occupied by $H_1$, and the resulting resource density must not exceed its initial value. Thus, resource regeneration is limited to restoring the initial landscape conditions, and resource densities cannot exceed their initial levels.

Shrub biomass does not regenerate during the simulated time interval. Thus, any biomass removed by $H_2$ remains depleted and produces a progressive reduction in refuge availability.

#### Mortality and refuge effects

Mortality of $H_1$ is associated with two mechanisms: predation and direct interactions with $H_2$. Local predation risk depends on shrub abundance, with greater vegetation cover providing more refuge.

Mortality events are activated according to the local vegetation coverage in each patch $(i,j)$, defined as

$$R_T(i,j) = R_1(i,j) + R_2(i,j) + R_3(i,j) + R_4(i,j). \tag{S10}$$

When $R_T$ falls below the threshold $R^*$, an $H_1$ individual may be removed from the population. If $H_2$ is absent from the selected patch, mortality can occur through predation with probability $e$. If $H_2$ is present, mortality associated with direct interactions occurs with probability $e'$. These interactions represent processes such as trampling or incidental mortality during grazing.

The stochastic mortality rule is given by

$$P(R^*, H_2) = \begin{cases} H_1(i,j) \xrightarrow{e} E_1(i,j) & R_T \le R^* \quad \& \quad H_2 = 0 \\ H_1(i,j)H_2(i,j) \xrightarrow{e'} E_1(i,j)H_2(i,j) & R_T \le R^* \quad \& \quad H_2 > 0 \\ 0 & R_t > R^* \end{cases} \tag{S11}$$

The parameter values used for $e$, $e'$, and $R^*$ are reported in Table S1.

##### Stochastic time evolution

The model evolves as a stochastic sequence of elementary updates. As in previous works [2, 3], we introduce stochastic variables $q_i$, which we refer to as classes, to ensure that all biological processes are treated as stochastic events. The occurrence of the biological processes considered in the model is associated with these classes. Thus, a patch is selected with probability $1/N$, and one of the following processes can then occur:

- With probability $q_1$, resource consumption occurs. The selected patch is first checked for the presence of individuals of $H_1$ and $H_2$ and for the availability of the corresponding resources. If both species are present, the order in which they attempt to consume is determined randomly. Each species then consumes sequentially according to its local occupancy density, with all individuals of the species present in the patch having the opportunity to consume. For each individual, at most one of the available resources is selected according to the corresponding consumption probabilities defined above. Thus, individuals of the same species may consume different resources within the same patch, while the total amount consumed depends on the number of individuals of each species present. Both species may consume during the same event, provided that the corresponding resources are available.

- With probability $q_2$, a mortality event affecting $H_1$ occurs. The selected patch must contain at least one $H_1$ individual for the event to take place. If the patch contains $H_2$, all $H_1$ individuals present in the patch are subject to mortality due to interactions with $H_2$, with probability $e'$. If $H_2$ is absent, mortality of each $H_1$ individual occurs through predation by predators present in the environment, with probability $e$.

- With probability $1 - q_1 - q_2$, a migration event occurs. The selected patch is checked for the presence of individuals of $H_1$ and $H_2$, and migration is evaluated for each species independently. For $H_1$, all individuals present in the selected patch have the opportunity to migrate, each with the corresponding migration probability. For $H_2$, migration is likewise evaluated if at least one individual is present.

  For each attempt at migration by an individual, a destination site is randomly selected among the neighboring sites that are eligible. For $H_1$, the destination site is selected within a radius $d$ (Eq. S3) from the current position, whereas for $H_2$, the destination is selected among first- and second-neighbor sites on the lattice. In both cases, the selected destination site must have sufficient capacity to accommodate the migrating individual. Once the destination site has been selected, migration occurs with the corresponding probability $m_i$, where for $H_1$ this probability also depends on the direction of movement relative to the higher-resource region, as described above. Thus, both species may migrate during the same migration event.

Once the rules governing the occurrence of the biological processes have been established, the temporal evolution of the system is implemented through a Monte Carlo algorithm. At each Monte Carlo step (MCS), a patch is randomly selected $N$ times. At each selection, one of the biological processes described above may occur, namely, resource consumption, migration of either herbivore population, or mortality of $H_1$. In this work, $q_1 = 0.75$, $q_2 = 0.08$ and $q_3 = 0.17$. The flowchart in Fig. S1 summarizes the sequence of the stochastic time evolution.

##### Temporal calibration

The number of Monte Carlo steps is linked to a biological time scale through the characteristic time required for $H_1$ to reach the higher-resource region A. This reference process is used to establish the correspondence between MCS and biological days. Once approximately 90% of the $H_1$ individuals have reached region A, the corresponding number of MCS is taken as the simulation time window. This interval is then divided by the total number of biological days to be represented, thereby defining the number of MCS corresponding to one biological day.

In this study, we consider a simulation period of 100 biological days, following the temporal windows adopted in previous studies [4]. The total simulation time ($t_{max}$) was set to $10^4$ MCS, corresponding to 100 MCS per biological day.

##### Consumption calibration

Resource consumption was calibrated using approximated daily consumption rates estimated from previous studies [4, 5]. Once the correspondence between MCS and biological days had been established, the consumption decrement $\Delta R_{m,n}$ for each resource $R_n$ consumed by species $m$ was

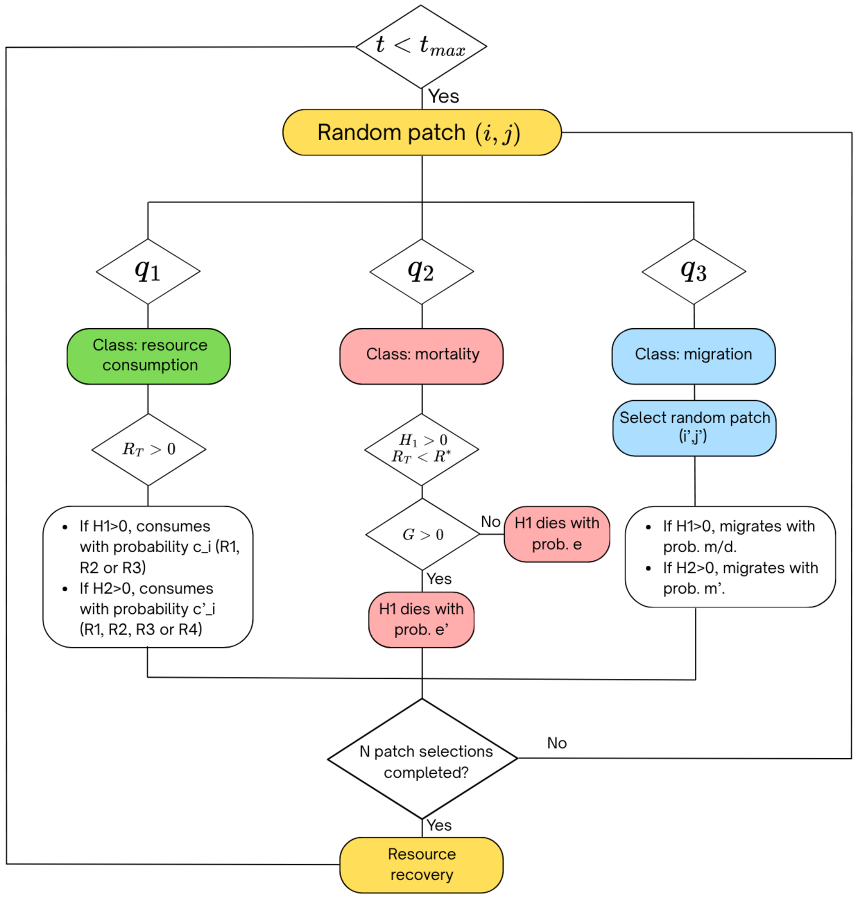


Figure S1: Flowchart of the individual-based stochastic model.

determined. Let $\rho_{m,n}$ denote the daily consumption rate of resource $R_n$ by species $m$, expressed in grams per individual per day, and let $s_{m,n}$ be the total number of successful feeding events involving that resource and species throughout the simulation. The average number of successful feeding events per individual per biological day was calculated as

$$\nu_{m,n} = \frac{s_{m,n}}{H_{m,0} t_d}, \tag{S12}$$

where $H_{m,0}$ is the initial density of the consumer population $m$, and $t_d$ is the total simulation time in biological days. The consumption decrement per successful feeding event was then obtained as

$$\Delta R_{m,n} = \frac{\rho_{m,n}}{\nu_{m,n}}. \tag{S13}$$

This calibration was performed at low initial densities for both $H_1$ and $H_2$, and the resulting values of $\Delta R_{m,n}$ were kept fixed as consumer densities increased. Values for $\rho_n$ and $\Delta R_{m,n}$ are presented in Table S1.

##### Resource recovery calibration

Resource recovery was implemented using recovery periods of 14 biological days for $R_1$ and $R_3$, and 21 biological days for $R_2$. These periods were converted into their corresponding numbers of MCS according to the established correspondence between simulation time and biological days. Resource recovery occurs gradually, through daily increments, and is bounded by the initial biomass of each resource. Thus, after each biological day, the biomass of resource $R_n$ in a patch $(i, j)$ can increase by an amount given by

$$\delta R_n = \frac{R_{0,n}}{\tau_n}, \tag{S14}$$

where $R_{0,n}$ is the initial biomass of resource $R_n$ in the patch and $\tau_n$ is its recovery period, expressed in biological days. This increment is added to the resource biomass in the patch, provided that the patch satisfies the regeneration conditions and the resulting biomass does not exceed its initial value.

#### Simulation experiments

The simulations were performed on landscapes containing $10,000$ patches. The initial number of $H_1$ individuals was varied between 0 and 150,000, and the initial number of $H_2$ individuals between 0 and 100. The resource carrying capacity $R$ was varied between 400 and 1000 $\mathrm{g, m^{-2}}$. Other model parameters were kept fixed at the values reported in Table S1, unless otherwise stated.

For each combination of control parameters, 5–15 independent realizations were performed for $10^4$ Monte Carlo steps. The number of realizations was adjusted according to the system's behavior and the statistical sampling required in each case. Initial conditions were independently generated for each realization according to the prescribed spatial distributions.

#### Parameterization

The model parameters used in the simulations are summarized in Table S1. The table reports the parameter values, their biological meaning, and the source or rationale used to define them. Some parameter values were obtained from empirical information when available, while others were set as model assumptions or varied systematically as control parameters. Unless otherwise stated, the values listed in the table were kept fixed across simulations.

Table S1: Model parameters used in the simulations. The table reports the parameter values, their description, and their source or rationale.

| Parameter | Description | Value(s) | Source |
|---|---|---|---|
| $L_x, L_y$ | Landscape dimensions | $100 \times 100$ patches | Model setup |
| $T$ | $H_1$ carrying capacity per patch | 60 individuals | Model setup |
| $G$ | $H_2$ carrying capacity per patch | 1 individual | Model setup |
| $R$ | Resource carrying capacity | [400-1000] $\mathrm{g\,m^{-2}}$ | Explored parameter |
| $H_{1,0}$ | Initial number of $H_1$ individuals | $[0 - 150000]$ | Explored parameter |
| $H_{2,0}$ | Number of $H_2$ individuals | $[0 - 70]$ | Explored parameter |
| $c_{1,1}$ | $H_1$ consumption prob. for $R_1$ | 0.52 | [5] |
| $c_{1,2}$ | $H_1$ consumption prob. for $R_2$ | 0.28 | [5] |
| $c_{1,3}$ | $H_1$ consumption prob. for $R_3$ | 0.20 | [5] |
| $c_{2,1}$ | $H_2$ consumption prob. for $R_1$ | 0.30 | [5] |
| $c_{2,2}$ | $H_2$ consumption prob. for $R_2$ | 0.30 | [5] |
| $c_{2,3}$ | $H_2$ consumption prob. for $R_3$ | 0.20 | [5] |
| $c_{2,4}$ | $H_2$ consumption prob. for shrubs | 0.20 | [5] |
| $e$ | $H_1$ mortality prob. due to predation | 0.0005 | Model setup |
| $e'$ | $H_1$ mortality prob. due to $H_2$ | 0.004 | Model setup |
| $m_0$ | $H_1$ migration prob. in A | 0.50 | Model setup |
| $m_1$ | $H_1$ migration prob. toward A | 1.00 | Model setup |
| $m_2$ | $H_1$ migration prob. away from A | 0.05 | Model setup |
| $m_3$ | $H_1$ migration prob. same distance from A | 0.20 | Model setup |
| $m'$ | $H_2$ migration prob. | 0.50 | Model setup |
| $\Delta R_{1,1}$ | Consumption decrement of $R_1$ by $H_1$ | 0.006 g | [5] |
| $\Delta R_{1,2}$ | Consumption decrement of $R_2$ by $H_1$ | 0.004 g | [5] |
| $\Delta R_{1,3}$ | Consumption decrement of $R_3$ by $H_1$ | 0.006 g | [5] |
| $\Delta R_{2,1}$ | Consumption decrement of $R_1$ by $H_2$ | 20 g | [1] |
| $\Delta R_{2,2}$ | Consumption decrement of $R_2$ by $H_2$ | 20 g | [1] |
| $\Delta R_{2,3}$ | Consumption decrement of $R_3$ by $H_2$ | 30 g | [1] |
| $\Delta R_{2,4}$ | Consumption decrement of $R_4$ by $H_2$ | 50 g | [1] |
| $f_A$ | Area A (% of landscape) | $\{20, 25, 30, 40, 43\}$ | Model setup |
| $f_B$ | Area B (% of landscape) | $\{80, 75, 70, 60, 57\}$ | Model setup |

### 2 Supplementary Results

#### Spatial dynamics at different stages of the simulation

Figure S2 shows the spatial distribution of the three renewable resources ($R_1$, $R_2$, and $R_3$) and the two herbivore populations at different times during the simulation. These figures complement those

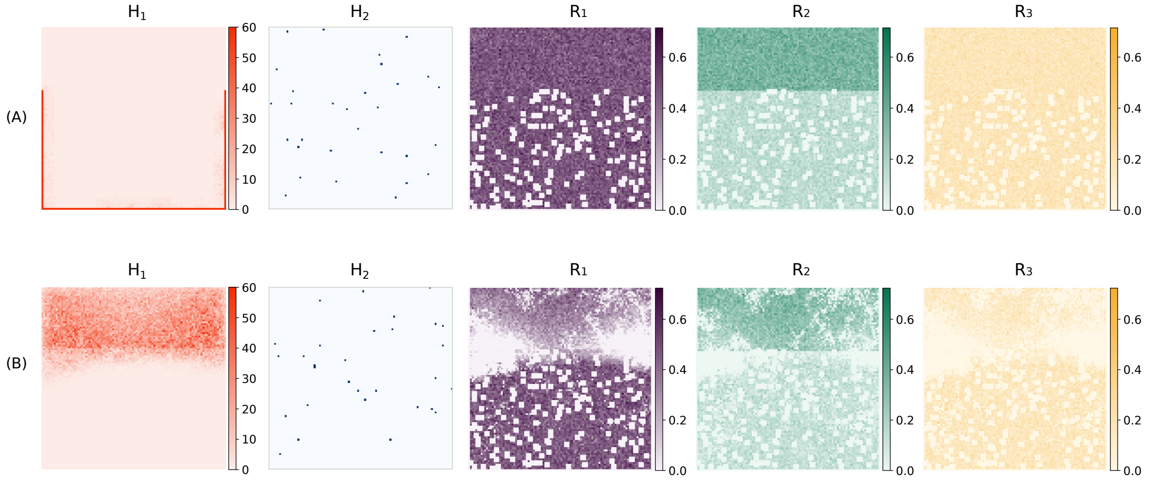


Figure S2: Spatial distributions of the three renewable vegetation resources ($R_1$, $R_2$, and $R_3$), $H_1$, and $H_2$ at different simulation times, for an initial $H_1$ density of 9 individuals m$^{-2}$, 30 individuals of $H_2$, and $R = 700$ g m$^{-2}$. (A) Initial configuration ($t = 0$). (B) Spatial distribution at an intermediate simulation time ($t = 50$ days).

shown in Fig. 2 A of the main text. At early times (panel A), corresponding to the initial stage of the dynamics, the spatial configuration reflects the imposed heterogeneity of the landscape. While $R_1$ and $R_3$ are randomly distributed across the landscape, $R_2$ is more abundant in area A than in area B. The distribution of $H_2$ extends across the landscape, whereas $H_1$ is initially confined to the edges of the grid. Shrubs ($R_4$) are not shown because they are consumed exclusively by $H_2$ and, although their depletion increases the mortality risk for $H_1$, they have a comparatively minor effect on the overall spatial dynamics.

Panel (B) shows the system at the midpoint of the simulation (5000 MCS $\equiv$ 50 biological days). At this stage, $H_1$ has begun to reach area A. The traces left by its movement across the landscape are visible as areas of resource depletion. Once $H_1$ reaches area A, individuals remain there and continue consuming resources. Additional snapshots of the dynamics are presented in the main paper.

#### Vegetation patch occupancy

Figure S3 shows the temporal dynamics of vegetation patch occupancy, expressed as the fraction of available patches occupied by each renewable resource. Panel A shows the dynamics of $R_1$, $R_2$, and $R_3$, together with the total biomass per patch, while panels B and C show the corresponding quantities restricted to areas A and B, respectively. In each case, occupancy fractions are normalized by the total number of available patches in the corresponding area. These results complement the biomass-based analysis presented in Fig. 1 of the main paper. The curves for $H_1$ are omitted from all panels to improve the visualization of the other quantities, as their different scales would otherwise hinder comparison.

The comparison between biomass and patch occupancy reveals that reductions in biomass occur before a substantial loss of occupied patches. Thus, vegetation can remain present in a large fraction of the landscape while its biomass has already been substantially reduced. This result highlights the difference between vegetation presence and the amount of forage available within occupied patches.

#### Remaining biomass varying R

Figure S4 shows the remaining biomass of each renewable resource after 100 days as a function of the initial $H_1$ density, for a fixed initial abundance of 30 individuals of $H_2$ and different values of the carrying capacity $R$ (400, 600 and 700). As shown in the main paper (Section 3b), the biomass of all three resources decreases as the initial $H_1$ density increases, although the magnitude of the response differs among resources and values of $R$. $R_2$ exhibits the strongest reduction, whereas $R_1$ and $R_3$ retain larger fractions of their initial biomass. The remaining biomass of $R_2$ is lower for smaller values of $R$ (400 g m$^{-2}$), and its decline with increasing initial $H_1$ density is more

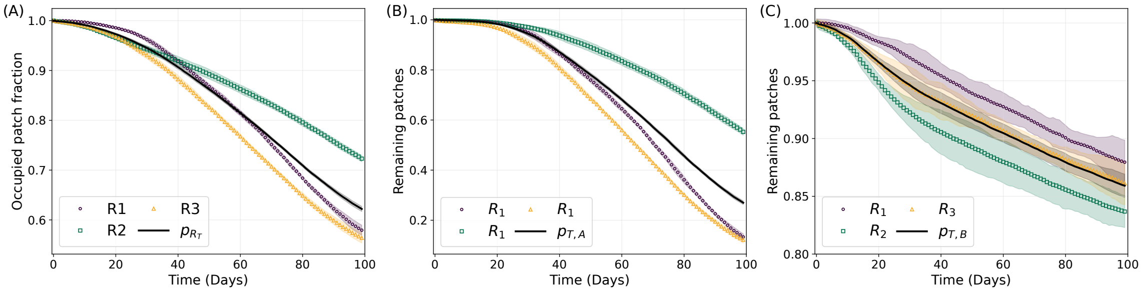


Figure S3: Temporal evolution of vegetation patch occupancy for each renewable resource and total renewable biomass ($R_\mathrm{T}$). (A) Entire landscape. (B) Area A. (C) Area B. Parameter values are as specified in the text, with an initial $H_1$ density of 9 individuals m$^{-2}$, 30 individuals of $H_2$, and $R = 700$ g m$^{-2}$. Each data point represents the average over five independent simulation runs, with shaded areas indicating the corresponding standard deviations. Different axis scales are used across panels to better highlight the results.

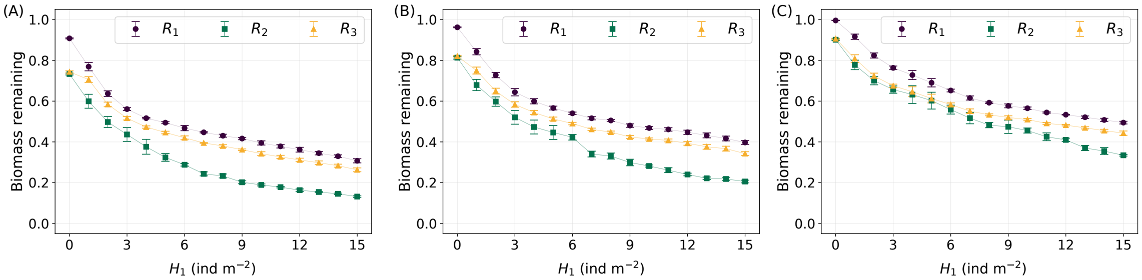


Figure S4: Biomass remaining after 100 days for each renewable resource as a function of initial $H_1$ density, for a fixed abundance of 30 individuals of $H_2$ and different values of $R$: (A) 400 g m$^{-2}$, (B) 600 g m$^{-2}$ and (C) 1000 g m$^{-2}$. Each data point represents the mean final biomass across five independent simulation runs. Final biomass is averaged over the last biological day (100 MCS). Error bars indicate the standard deviation across the five runs. Final biomass is normalized by the initial biomass of the corresponding resource.

pronounced. In contrast, for larger values of $R$ (1000 g m$^{-2}$), $R_2$ retains a higher biomass and exhibits a less pronounced decline as the initial $H_1$ density increases.

For each independent simulation run, the final biomass was calculated as the average over the last 100 MCS, corresponding to the final biological day. The mean final biomass was then calculated across the five independent runs. Error bars indicate the standard deviations of the final biomass values.

#### Varying landscape structure

To investigate the effect of the relative sizes of areas A and B, we varied the fraction of the landscape occupied by each area while keeping the local biomass of resource $R_2$ per patch constant in each area. Here, $r_A$ and $r_B$ denote the biomass of $R_2$ per patch in areas A and B, respectively, and $k = r_A/r_B$ quantifies the relative resource availability between the two areas. Thus, both $r_A$ and $r_B$, as well as their ratio $k$, were kept constant throughout the analysis. Consequently, the total initial biomass of $R_2$ varied with the relative sizes of the two areas. The values of $k$, $r_A$, and $r_B$ were set to those corresponding to the reference configuration used in the previous results, with $R = 700$ g m$^{-2}$, an initial $H_1$ density of 10 individuals m$^{-2}$ and 30 individuals of $H_2$. Thus the area ratio $f_A/f_B = 0.54$, corresponding to 35% of the landscape occupied by area A and 65% by area B. Accordingly, $k = 2.36$, $r_A \approx 250$ g per patch, and $r_B \approx 105$ g per patch.

Figure S5 shows the initial and final biomass of $R_2$ as a function of $f_A/f_B$. The initial biomass increases with the extent of region A because $R_2$ is preferentially distributed in this region, while its local biomass is kept unchanged across landscape configurations. In contrast, the final biomass remains approximately constant despite the increase in initial resource availability. Thus, a larger initial biomass does not translate into a correspondingly larger biomass remaining after 100 days. This pattern indicates greater net resource depletion as the productive region expands, although the final biomass also reflects resource regeneration during the simulation.

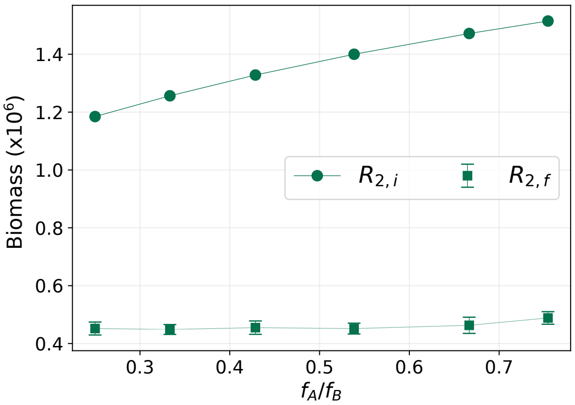


Figure S5: Initial and remaining biomass of resource $R_2$ as a function of the fraction $f_A/f_B$, for a initial density of 10 individual m$^{-}$2 for $H_1$, 30 individuals for $H_2$ and $R = 700$ g m$^{-}$2. Each data point represents the final biomass, calculated by averaging over the last 100 MCS of each simulation run, as described above, and then averaged across five independent simulation runs of $10^4$ MCS. Error bars indicate the corresponding standard deviations.

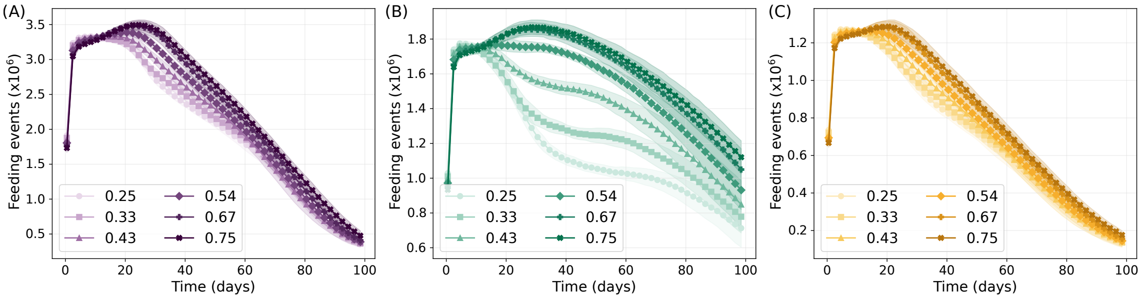


Figure S6: Temporal evolution of feeding events for the three renewable resources: (A) $R_1$, (B) $R_2$, and (C) $R_3$. Parameters: initial $H_1$ density of 10 individuals m$^{-2}$, 30 individuals of $H_2$, and $R = 700$ g m$^{-2}$. Each data point represents the total number of successful feeding events per day, averaged over 15 independent simulation runs. The shaded areas indicate the corresponding standard deviations.

Figure S6 shows the temporal evolution of successful feeding events involving each of the three renewable resources for different values of the area ratio $f_A/f_B$, which represents the relative sizes of regions A and B. These events contribute to the resource consumption dynamics underlying the total renewable-resource biomass shown in Fig. 3B of the main paper. $R_2$ exhibits the strongest response to changes in the area ratio: as $f_A/f_B$ increases, the number of successful feeding events involving $R_2$ increases, reflecting its greater availability in region A. In contrast, feeding events involving $R_1$ and $R_3$ show smaller differences among landscape configurations. The increase in feeding activity involving $R_2$ is consistent with the greater net depletion of this resource despite its increasing initial biomass (Fig. S5). Together, these results suggest that expanding region A increases resource availability while also sustaining feeding activity that counteracts the effect of this additional availability on the final biomass.
